## Supplementary material for "Relating simulation studies by provenance—Developing a family of Wnt signaling models": S1 Appendix

February 15, 2021

### 1 List of names and references of studies shown in provenance graphs

This list shows the names used in the provenance graphs as well as the corresponding references.

**Wnt simulation studies:**

- Chen et al. 2014 [1]
- Cho et al. 2006 [2]
- Goldbeter and Pourquié 2008 [3]
- Haack et al. 2015 [4]
- Haack et al. 2020 [5]
- Kim et al. 2007 [6]
- Kogan et al. 2012 [7]
- Krüger and Heinrich 2004 [8]
- Lee et al. 2001 [9]
- Mazemondet et al. 2012 [10]
- Mirams et al. 2010 [11]
- Padala et al. 2017 [12]
- Rodriguez et al. 2007 [13]
- Sick et al. 2006 [14]
- Staehlke et al. 2020 [15]
- van Leeuwen et al. 2007 [16]
- van Leeuwen et al. 2009 [17]
- Wang et al. 2013 [18]
- Wawra et al. 2007 [19]

**Additionally used studies:**

- Bafico et al. 2001 [20]
- Bernard et al. 2006 [21]
- Bourhis et al. 2010 [22]
- Brown et al. 2004 [23]
- Cho et al. 2003 [24]
- Dajani et al. 2003 [25]
- Dequeant et al. 2006 [26]
- Galli et al. 2006 [27]
- Goh and Sorkin 2013 [28]
- Hannoush 2008 [29]
- Hernandez et al. 2012 [30]
- Hirata et al. 2004 [31]
- Howard et al. 2001 [32]
- Jho et al. 2002 [33]
- Kim et al. 2006 [34]
- Kim et al. 2013 [35]
- Krieghoff et al. 2006 [36]
- Lee et al. 2001 [9]
- Lewis 2003 [37]
- Mazemondet et al. 2011 [38]
- Meineke et al. 2001 [39]
- Monk 2003 [40]
- Orton et al. 2009 [41]
- Pralle et al. 2000 [42]
- Prior et al. 2003 [43]
- Rida et al. 2004 [44]
- Sakane et al. 2010 [45]
- Salic et al. 2000 [46]
- Swat et al. 2004 [47]
- Wawrzak et al. 2007 [48]
- Yamamoto et al. 2006 [49]
- Yamamoto et al. 2008 [50]
