## Supplementary figures and images for "Relating simulation studies by provenance—Developing a family of Wnt signaling models"

### S1 Figure

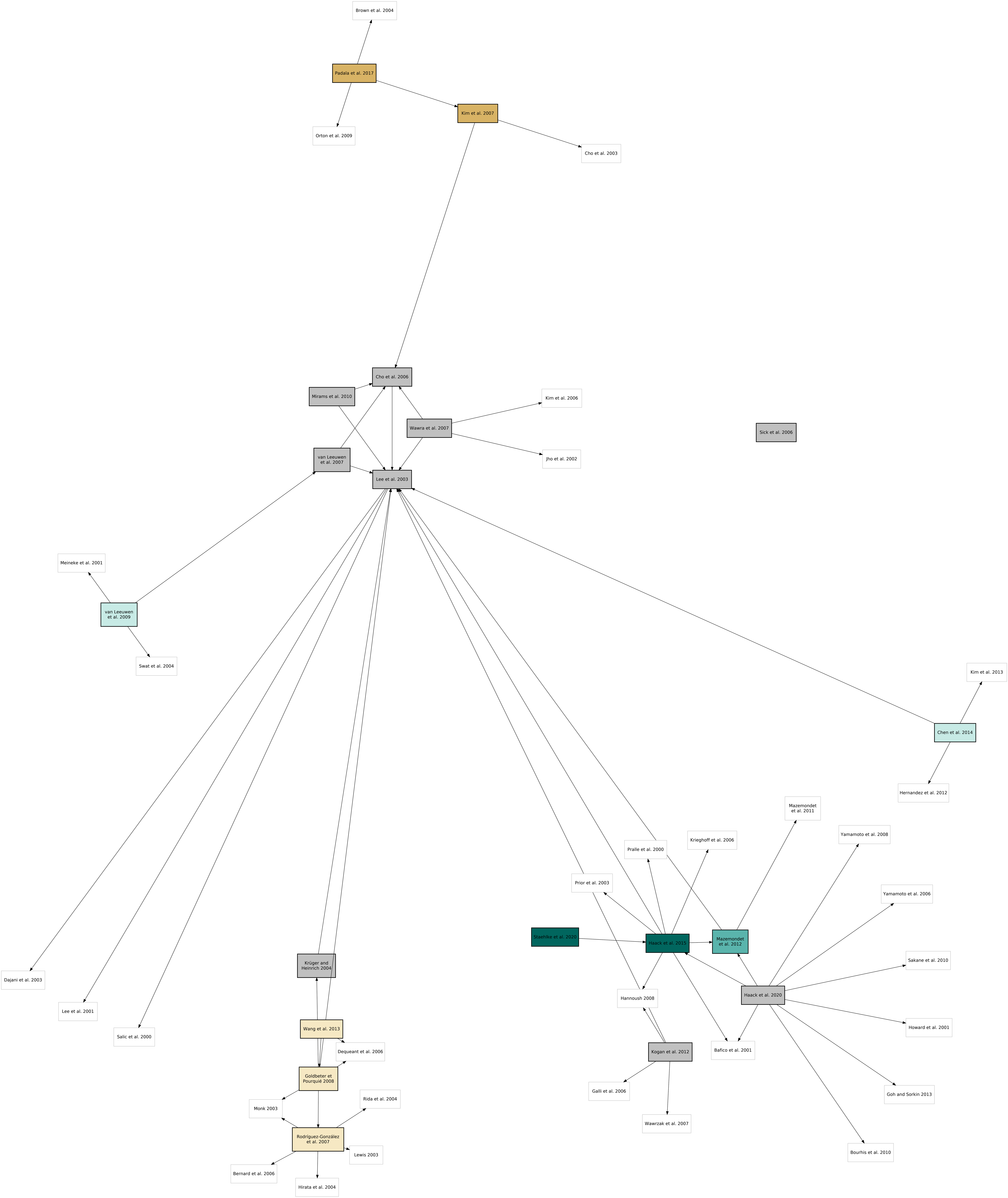
